# Targeted loss of heterozygosity in *Candida albicans* using CRISPR-Cas9

**DOI:** 10.1101/2025.05.13.653461

**Authors:** Philippe C Després, Nicholas C Gervais, Meea Fogal, Ruby KJ Rogers, Christina A Cuomo, Rebecca S Shapiro

## Abstract

The diploid genome of the fungal pathogen *Candida albicans* is highly heterozygous, with most allele pairs diverging at either the coding or regulatory level. When faced with selection pressure like antifungal exposure, this hidden genetic diversity can provide a reservoir of adaptive mutations through loss of heterozygosity (LOH) events. Validating the potential phenotypic impact of LOH events observed in clinical or experimentally evolved strains can be difficult due to the challenge of precisely targeting one allele over the other. Here, we show that a CRISPR-Cas9 system can be used to overcome this challenge. By designing allele-specific guide RNA sequences, we can induce targeted, directed LOH events, which we validate by whole-genome long-read sequencing. Using this approach, we efficiently recapitulate a recently described LOH event that increases resistance to the antifungal fluconazole. Additionally, we find that the recombination tracts of these induced LOH events have similar lengths to those observed naturally. To facilitate future use of this method, we provide a database of allele-specific sgRNA sequences for Cas9 that provide near genome-wide coverage of heterozygous sites through either direct or indirect targeting. This approach will be useful in probing the adaptive role of LOH events in this important human pathogen.

## Introduction

*Candida albicans* is a common member of the human microbiota and an opportunistic pathogen responsible for hundreds of thousands of life-threatening infections annually (Denning 2024). Unlike other diploid yeasts such as *Saccharomyces cerevisiae*, *C. albicans* is thought to rarely undergo sexual reproduction, propagating mostly clonally (Bedekovic and Usher 2023). Recent genomic analysis suggests that the ancestor of *C. albicans* was a hybrid between two lineages with ∼3% nucleotide divergence that lost the ability to efficiently undergo sexual reproduction shortly after hybridization (Mixão and Gabaldón 2020). While many of these heterozygous sites were lost during the course of evolution, tens of thousands persist or have arisen across the *C. albicans* genome. These divergent alleles can differ in terms of expression, translation, and function (Muzzey *et al*. 2014; Liang and Bennett 2019). Naturally occurring loss of heterozygosity (LOH) events can lead to one variant becoming homozygous and the other being lost. Lineage-specific LOHs are one of the major evolutionary mechanisms driving divergence between *C. albicans* populations (Wang *et al*. 2018).

As heterozygous sites can serve as reservoirs of adaptive variants in the face of strong selective pressure, LOH events are also thought to play an important role in the evolution of antifungal drug resistance (Ford *et al*. 2015). As an example, *de novo* mutations in the *TAC1* and *MRR1* transcription factors can become homozygous through LOH events, resulting in a dramatic increase in drug resistance (Coste *et al*. 2006, 2007; Dunkel *et al*. 2008). Additionally, recessive or incompletely dominant loss-of-function (LOF) alleles that are buffered by the presence of a functional copy on the homologous chromosome can become homozygous and have dramatic effects on phenotype. *C. albicans* Clade I isolates are known to possess higher intrinsic resistance to the antifungal 5-fluorocytosine due to a heterozygous LOF mutation in *FUR1* (Dodgson *et al*. 2004). When a LOH event makes this mutation homozygous, it results in complete resistance to the drug. Recently, a laboratory evolution experiment demonstrated that LOH events in *KSR1* can result in increased resistance to the antifungal fluconazole, decreasing susceptibility by 500-fold when combined with a segmental amplification on chromosome 4 (Vande Zande *et al*. 2024). Some evidence suggests that cellular stresses linked with the host environment and antifungal treatment could increase the rate of LOH in *C. albicans* (Forche *et al*. 2011; Popp *et al*. 2019), increasing the rate of evolution via this type of event.

Despite the importance of this class of mutational events, the complexities of reproducing LOH using genetic manipulation have often prevented in-depth study and validation of their role in the evolution of new phenotypes. Traditional approaches that rely on homologous recombination cannot be specifically targeted towards one allele or the other. This leads to more complex design strategies, which can make the strain construction process arduous and requires many rounds of transformations, which increase the chance of off-target mutations and structural rearrangements (Marton *et al*. 2020). Recent developments in CRISPR-Cas9-mediated genome editing have allowed for faster and more streamlined strain engineering strategies in *C. albicans* (Vyas *et al*. 2015; Min *et al*. 2016; Nguyen *et al*. 2017; Shapiro *et al*. 2017; Uthayakumar *et al*. 2020), but no method specifically designed to generate LOH events has been described yet. Here, we show that allele-specific CRISPR single guide RNAs (sgRNAs) can be used to efficiently induce LOH events in *C. albicans*. Using long-read sequencing, we show that the associated recombination tracts are comparable to naturally occurring LOH events and that off-target effects are rare. Finally, we validate our approach using a recently identified LOH involved in azole resistance and provide computational tools for allele-specific sgRNA identification to perform similar experiments. This approach will facilitate direct work with this understudied mutation class in *C. albicans*.

## Results

The Cas9-sgRNA complex recognizes a 20-nucleotide sequence followed by a protospacer adjacent motif (PAM) that must match the nucleotide pattern ‘NGG’. This PAM sequence is essential for both Cas9 binding and DNA cleavage (Nishimasu *et al*. 2014), meaning that any disruption of the motif could abolish Cas9 activity. Based on this, we hypothesized that we could use CRISPR-Cas9 to engineer targeted LOH events in *C. albicans* by exploiting sgRNAs where an intact downstream PAM motif is found exclusively in one of the alleles. Alternatively, mismatches in the sgRNA target sequence can abolish Cas9 binding and activity, especially if they occur within the last 12 bases (often referred to as the ‘seed’ sequence). Both scenarios should allow for specific cleavage of only one of the alleles until the DSB is repaired using the non-targeted allele as a template, yielding a controlled LOH (Figure 1A).

**Figure 1:**
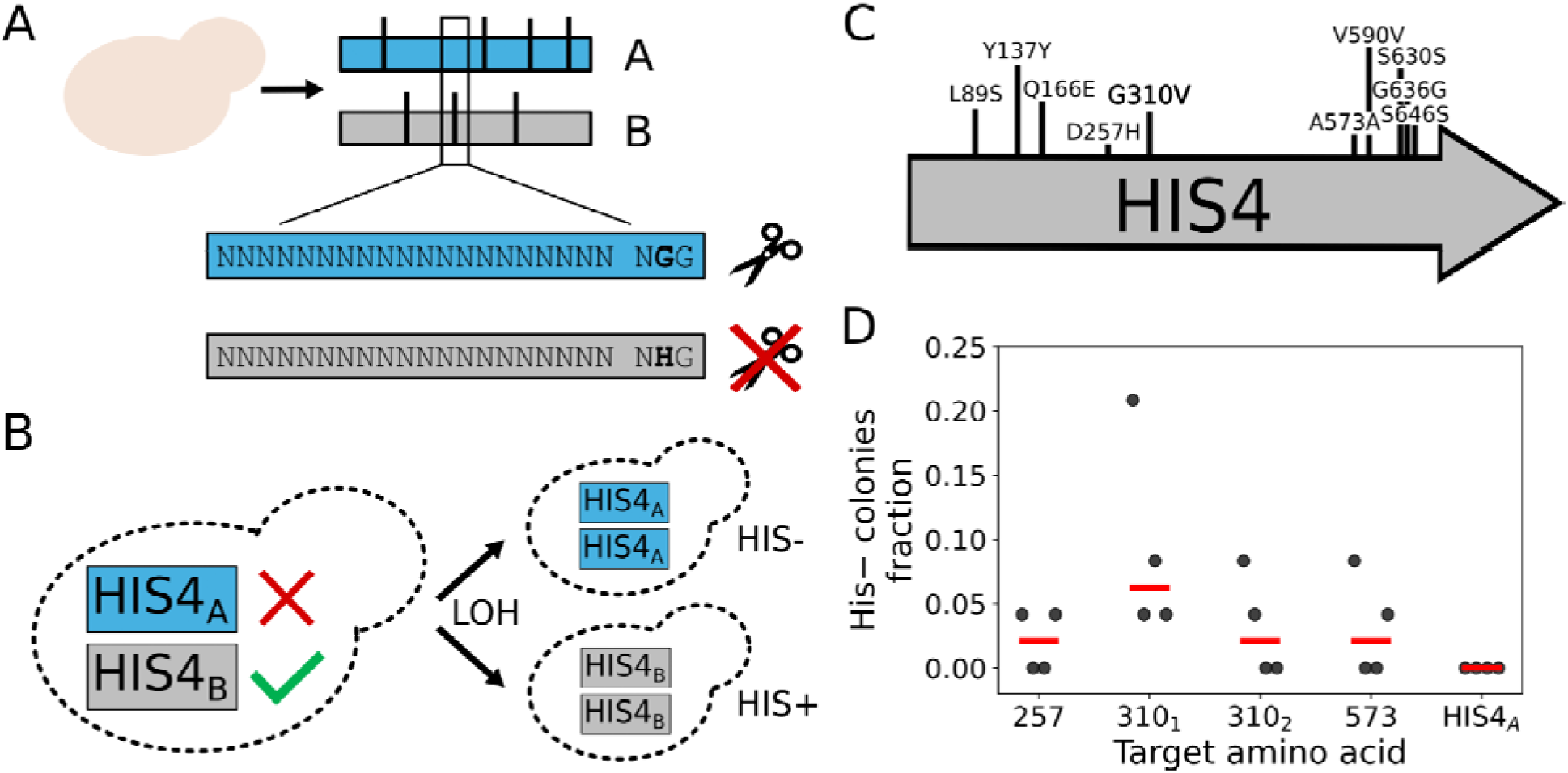
A method for targeted LOH engineering in *Candida albicans*. **A)** Heterozygous single-nucleotide polymorphisms (SNPs) within the *C. albicans* genome that coincide with potential sgRNA target sites allow allele-specific targeting of CRISPR-Cas9 mediated double-stranded breaks. **B)** A heterozygous loss-of-function mutation in *HIS4* present in SC5314 makes it an ideal test locus for LOH engineering. Conversion of *HIS4_B_* to *HIS4_A_* results in histidine auxotrophy (HIS-), which can be detected by spot assay on histidine drop-out media. **C)** Heterozygous sites in *HIS4* (allele A is shown first). Both missense and synonymous variants are depicted. **D)** HIS-conversion rate of sgRNAs targeting a subset of *HIS4* heterozygous sites. The causal mutation associated with *HIS4_A_* LOF, G310V, is shown in bold. Each transformation was performed in quadruplicate. The control sgRNA targeting *HIS4_A_*also targets position 310.

To test this approach, we used the *HIS4* locus as a model (Altboum *et al*. 1990). In the *C. albicans* SC5314 strain (Fonzi and Irwin 1993), the B allele of *HIS4* is functional, while the A allele is inactive due to the missense mutation Gly310Val (Gómez-Raja *et al*. 2008). If LOH is inducible via CRISPR-Cas9, targeting the wild-type B allele with the CRISPR system should render the non-functional A allele homozygous, resulting in histidine auxotrophy (Figure 1B). Conversely, targeting the A allele should not result in phenotypic changes if LOH occurs, allowing us to get a blind estimate of the CRISPR editing rate. We designed four B allele-specific sgRNAs: two directly targeting Gly310, and two targeting heterozygous variants located in proximity to Gly310 (D257H and A573A) (Figure 1C). As a control, we also included an additional A allele-specific sgRNA targeting Val310. The sgRNAs were cloned into an integrative CRISPR-Cas9 editing vector that we previously optimized (Bédard *et al*. 2024b), and transformed into *C. albicans*. Transformants were then transferred to media with or without histidine to test for the auxotrophy. All *HIS4_B_* targeting sgRNAs produced histidine auxotrophs at various rates, while the control *HIS4_A_* targeting sgRNA did not. The two most efficient sgRNAs directly targeted Gly310, suggesting that direct targeting of sites of interest is an efficient approach for LOH generation.

We investigated whether the histidine auxotrophy phenotypes we observed were due to the expected LOH, as opposed to the result of indels or missense mutations introduced by DNA repair processes. Previous work in *C. albicans* (Vyas et al. 2015) suggests that, similar to what has been observed in *Saccharomyces cerevisiae* (DiCarlo *et al*. 2013), CRISPR-Cas9 editing without a donor template doesn’t usually result in significant on-target editing. This suggests that most auxotrophy phenotypes should be the result of on-target recombination with the non-targeted allele. Nevertheless, to validate that auxotrophic strains harbored an LOH edit, we genotyped the *HIS4* locus by Sanger sequencing for a subset of strains generated using the sgRNA yielding the highest frequency of auxotrophs. We found that all auxotrophs had become homozygous for the G310V mutation, demonstrating that our approach was successful at inducing LOH.

While the results of Sanger sequencing indicated successful gene conversion, they did not allow us to precisely identify LOH breakpoints located further away from the DSB site. To address this and examine potential off-target effects, we obtained whole-genome data for four independent *HIS4_B_* strains using long-read sequencing. The depth of sequencing coverage was sufficient to call homozygous and heterozygous sites and accurately map breakpoints around the cut site (Table S1). For all strains, whole-genome sequencing data validated the presence of LOH events at the *HIS4* locus as determined previously (Figure 2). The recombination tracts generally spanned more than 1 kb (minimum = 0.66 kb bp, max = 15 kb), explaining why Sanger had been insufficient for precise mapping. However, not all events had clear boundaries: for some samples, the A allele frequency graph showed two distinct plateaus, suggesting multiple recombination outcomes were present in the same sample. We examined the long reads spanning both the target site and the secondary junction and found that variants in the extended plateau were always physically linked with one another and not distributed randomly between reads. Additionally, read coverage distributions did not show any increases supporting chromosome IV aneuploidy as the explanation for these two distinct edit outcomes within the same sequencing library. Instead, this suggests that colonies isolated after LOH editing can have heterogeneous genotypes, which can carry over to downstream experiments. In addition, we also tested the effect of providing a recombination template for LOH as donor DNA during the CRISPR transformation. We found that this increased the editing rate as measured by histidine auxotrophy, providing a potential avenue to increase efficiency (Figure S1). Whole-genome sequencing thus validated the targeted CRISPR-Cas9-induced LOH events and allowed for precise mapping of the resulting edits, even in mixed samples.

**Figure 2:**
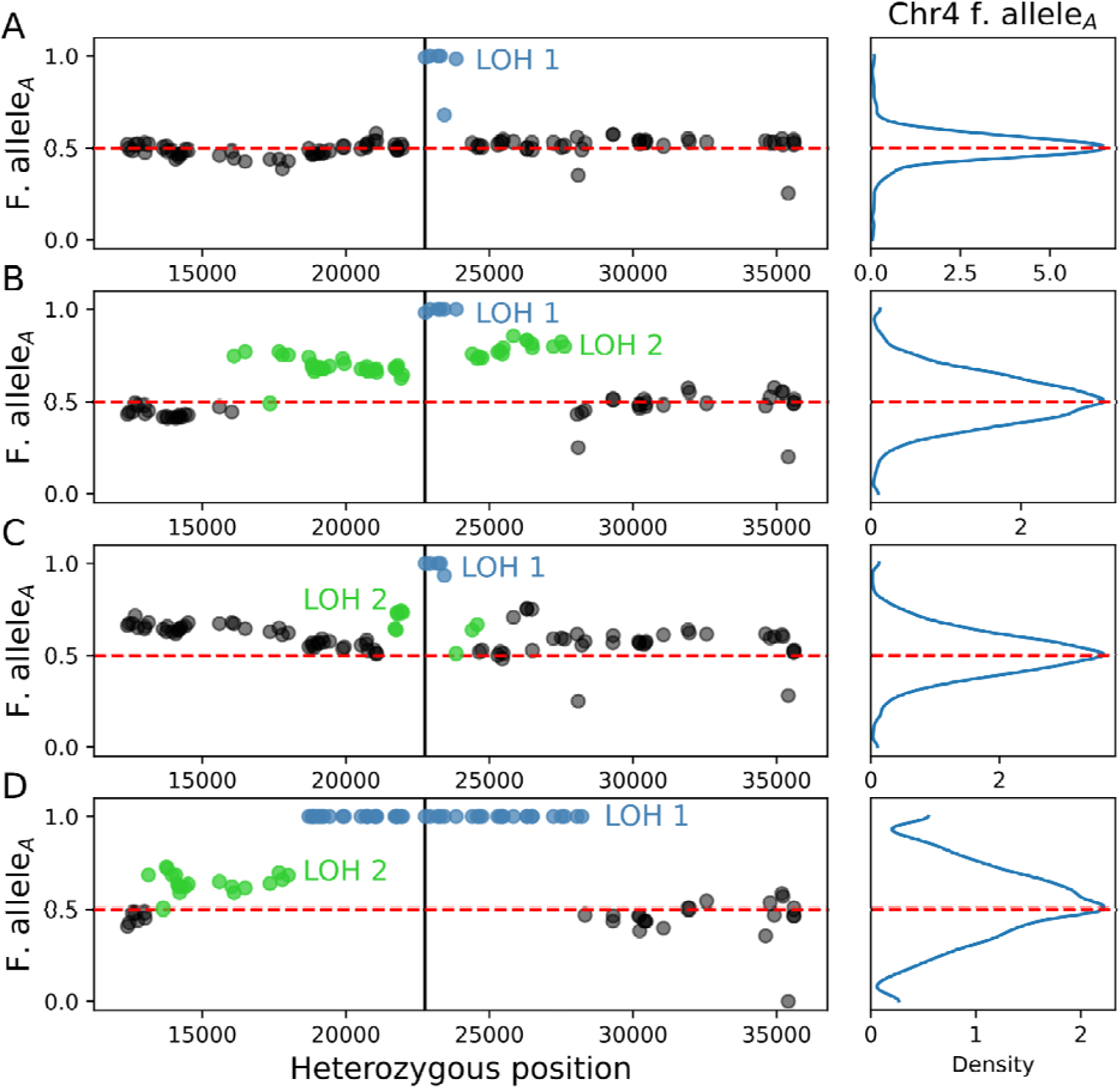
CRISPR-Cas9 induces short tract LOH events resulting in histidine auxotrophy. Chromosome A allele frequency around the *HIS4* heterozygous SNP targeted by CRISPR-Cas9 for auxotrophic strains **A)** HIS4_AA_1, **B)** HIS4_AA_2, **C)** HIS4_AA_3 and **D)** HIS4_AA_4. The vertical black line shows the position of the causal SNP, while the red dotted line shows the expected allele frequency of 0.5, and LOH tracts are colored according to the different events inferred from the sequencing data. The density plots on the right show the overall distribution of chromosome A allele frequencies across the full length of chromosome 4.

Genome editing approaches require a minimum level of efficiency to be scalable. Not all LOH events might result in an obvious or selectable phenotype on which to base clone validation. Unlike conversion to *HIS4_A_*/*HIS4_A_*, (Figure 1), conversion to *HIS4_B_*/*HIS4_B_*does not result in *HIS4* inactivation, and thus does not result in any discernible phenotype compared to the wild-type meaning that colonies with successful LOH events cannot be readily screened. To get another estimate of the efficiency rate of our approach, we also genotyped five clones generated with the control *HIS4_A_* sgRNA and were able to successfully identify three clones that appeared to bear *HIS4_A_* to *HIS4_B_* LOH events. Because editing appears to often result in heterogeneous colonies, it is possible that our previous experiment underestimated the actual editing rate. As colonies were resuspended and replica spotted at high cell density to test for histidine auxotrophy, any colonies with a mix of edited and wild-type genotypes would result in spot growth, camouflaging the presence of the edited subpopulation. This could explain the higher-than-expected rate of conversion in the *HIS4_B_*/*HIS4_B_* strains we genotyped.

Next, we validated the three *HIS4_A_* to *HIS4_B_* edited strains using long-read genome sequencing to confirm LOH events and the desired conversion to the *HIS4_B_* allele, as well as an unedited strain as a control (Figure 3). One of the edited strains had alterations in ploidy as determined by relative coverage analysis (Table S1), an outcome that has already been reported after both standard and CRISPR transformations in *C. albicans* (Marton *et al*. 2020). These results suggest that LOH induction is efficient enough to be applied without direct selection for a phenotype of interest, especially if a donor DNA template is provided, allowing for the use of our method to study a large diversity of LOH events in *C. albicans*.

**Figure 3:**
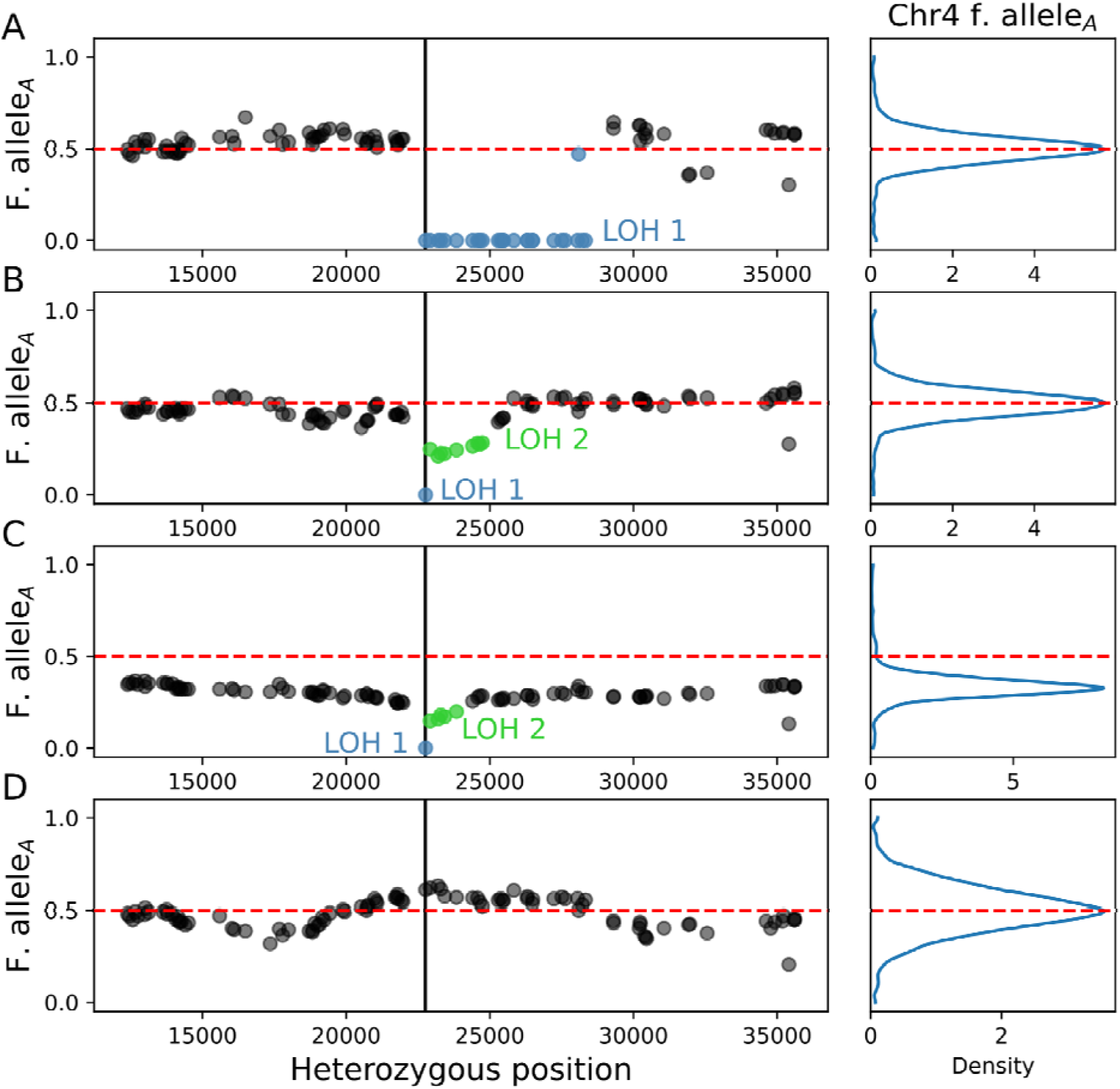
CRISPR-Cas9 induced LOHs with *HIS4_A_* targeting sgRNAs. Chromosome A allele frequency around the *HIS4* heterozygous SNP targeted by CRISPR-Cas9 for auxotrophic strains **A)** BB_1, **B)** BB_2, **C)** BB_3 (aneuploid for chromosomes 4 and 5), and **D)** BB_4 (no LOH detected). The vertical black line shows the position of the causal SNP, while the red dotted line shows the expected allele frequency of 0.5, and LOH tracts are colored according to the different events inferred from the sequencing data. The density plots on the right show the overall distribution of chromosome A allele frequencies across the full length of chromosome 4.

Structural variants like LOH events are often reported in antifungal-resistant isolate genomes of *C. albicans* and other fungal pathogens (White 1997; Coste *et al*. 2006; Dunkel *et al*. 2008; Heilmann *et al*. 2010; Ford *et al*. 2015; Ene *et al*. 2018), but their contributions are seldom validated experimentally. To test if our approach could be applied to studying LOH events associated with antifungal drug resistance, we attempted to reproduce recent results that showed that partial LOH of *KSR1_B_* into *KSR1_A_* was linked with increased fluconazole resistance in experimentally evolved isolates (Vande Zande *et al*. 2024). We designed sgRNAs targeting the causal heterozygous site that was previously identified in *KRS1* (Figure 4A). After transformation, a few colonies were picked randomly from plates and assayed for fluconazole resistance via minimum inhibitory concentration (MIC) assays. Across multiple independent experiments, we found that a median of 33% of transformants showed increased resistance to fluconazole when using the best-performing sgRNA (plasmid Cas9-KSR1_A-2, Figure 4B). We used long-read sequencing to genotype four strains with elevated MICs from different transformation reactions. In three out of four cases, we could validate the presence of LOH events with short recombination tracts at the target site (Figure 4C-E). One of the positive samples had increased A allele frequency across its entire genome, suggesting one of the subclones might have become homozygous for the A copy of chromosome R. Finally, the resistant strain in which we could not validate the intended LOH had a more complex rearrangement involving multiple LOH and a point mutation at the target site, as well as trisomy of chromosome 5 (Table S1, Figure S2). Despite the higher occurrence of rearrangements, our approach was still effective in reproducing an LOH event directly involved in the emergence of antifungal resistance.

**Figure 4:**
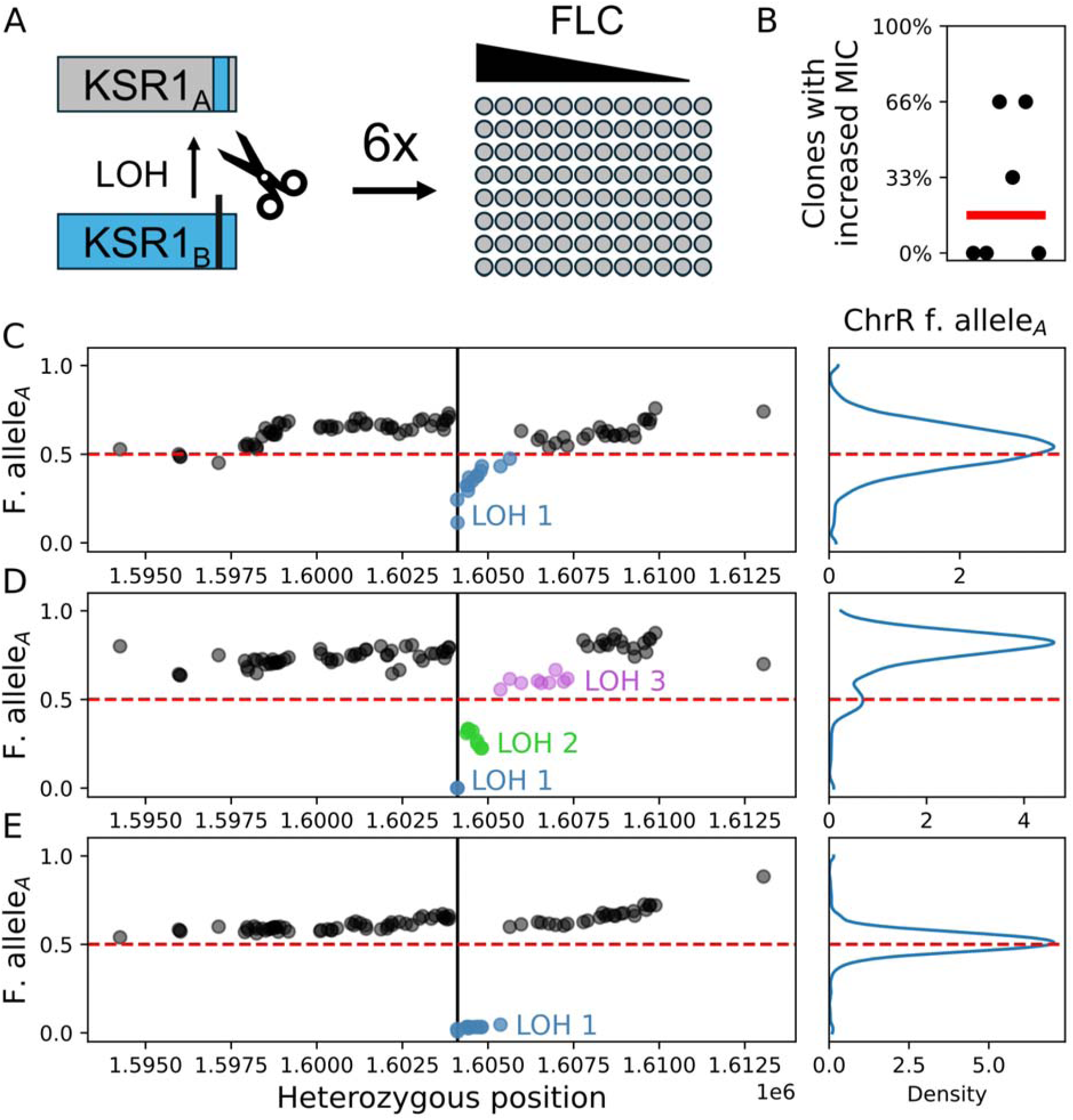
An LOH with known effects on resistance can be reproduced via genome editing. **A)** The LOH event in *KSR1* that is associated with increased fluconazole resistance was engineered by transforming cells with a CRISPR-Cas9 vector bearing a *KSR1_A_* targeting sgRNA (Cas9-KSR1_A-2) in multiple independent reactions. Three transformants per plate were picked and assayed for increased MICs. **B)** Fraction of transformants with increased fluconazole MIC per plate. The red line represents the median percentage of clones with increased resistance across plates. **C)**, **D)** and **E)** Chromosome A allele frequency around the *HIS4* heterozygous SNP targeted by CRISPR-Cas9 for auxotrophic strains KSR1_2, KSR1_3 and KSR1_4, respectively. The vertical black line shows the position of the causal SNP, while the red dotted line shows the expected allele frequency of 0.5, and LOH tracts are colored according to the different events inferred from the sequencing data. The density plots on the right show the overall distribution of chromosome A allele frequencies across the full length of chromosome R.

Here, we have demonstrated that targeted, allele-specific LOH induction is possible in *C. albicans.* Nonetheless, the broad applicability of this technique will depend on how well the induced events approximate those found in the wild, as well as how many heterozygous sites are amenable to allele-specific CRISPR targeting. Across all CRISPR-induced LOH events that we monitored by whole-genome sequencing, the median recombination tract length was 1.264 kb (Figure 5A, Table S2), which is close to what has been observed for LOH events in passages strains (Ene *et al*. 2018; Liang and Bennett 2019). Additionally, for both *HIS4 and KSR1*, we could identify recombination tracts spanning one nucleotide within the heterogeneous population, suggesting that clones with “perfect” edits could readily be isolated by subculturing from transformant colonies.

**Figure 5:**
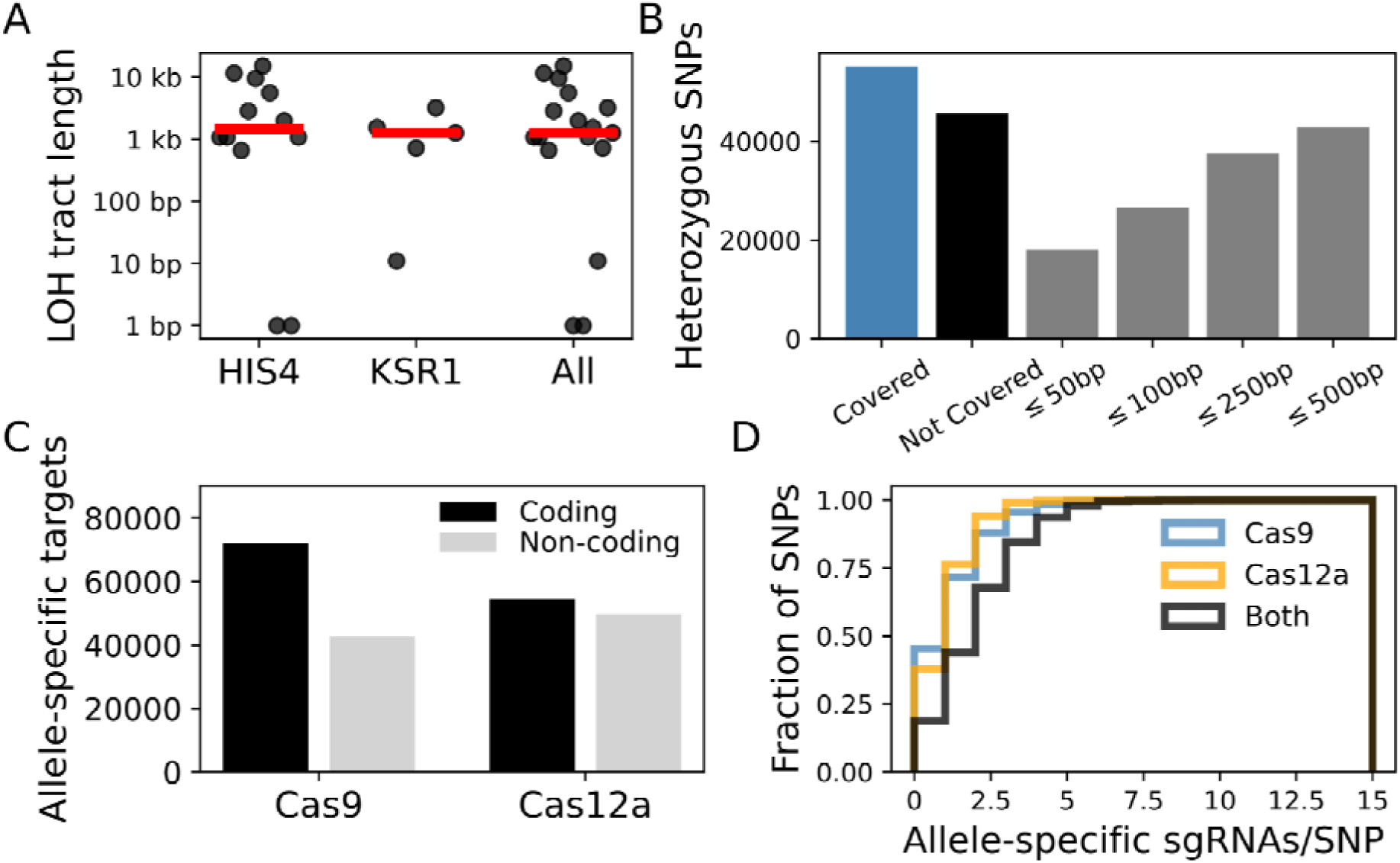
Genome-scale applicability of CRISPR-LOH in C. albicans. **A)** Recombination tract length for induced LOH events in *HIS4, KSR1,* and for both genes. Tract length was calculated as the length of the genomic region with increased or decreased A allele frequency. **B)** Allele-specific sgRNA coverage of heterozygous SNPs. Distance to the next heterozygous site was measured in both directions. **C)** Distribution of Cas9 and Cas12a allele-specific targets in coding and non-coding regions. **D)** Cumulative density of allele-specific targets per heterozygous SNPs (n=56338) in C. albicans for Cas9, Cas12a, and their combination.

Next, we wanted to test if this approach was broadly applicable at the genome scale. Using the SC5314 phased diploid reference genome (Skrzypek *et al*. 2017), we identified all potential Cas9 sgRNA target sites specific to one homologous chromosome or the other. We found that allele-specific sgRNAs were frequent, with a median density of ∼3 sgRNAs/kb depending on the level of heterozygosity of the chromosome (Figure S3). This is similar to the average density of heterozygous sites, which is estimated to be 1/200-300 bp (Liang and Bennett 2019). This high sgRNA density results in ∼55% direct coverage of heterozygous SNPs across the diploid genome (Figure 5B), either through allele specific PAMs or sgRNA sequences. As our approach usually generates recombination tracts spanning ∼1-3 kb, we also asked what fraction of sites not directly covered could be indirectly targeted via another nearby heterozygous site. We found that in over 90% of cases, another target amenable to editing is present within a 500 bp window. By combining both direct and indirect targeting, almost all heterozygous sites in the *C. albicans* genome can be targeted for LOH engineering.

Since the emergence of CRISPR-based genome editing systems, the repertoire of available targetable nucleases has greatly expanded. Many of these enzymes have different sgRNA and PAM requirements compared to Cas9, which can further increase the proportion of the genome amenable to direct targeting for genome editing. This is particularly interesting in the context of *C. albicans* and some of its close relatives, whose genomes have a relatively low GC content (Lynch *et al*. 2010), which can make the NGG PAM more restrictive. In addition, intergenic regions are often more AT-rich, leading to lower target density. One of the most common alternatives to Cas9 is Cas12a, which instead requires a TTTV PAM motif (Zetsche *et al*. 2015). While it has not yet been adapted for use in *C. albicans*, we also generated a database of allele-specific target sequences. As expected, Cas12a had better relative and absolute coverage in intergenic regions compared to Cas9 (Figure 5C). Combining the two sets of targets led to improved coverage for direct targeting (Figure 5D). The resulting database of sgRNA sequences is provided as Table S5 and Table S6 for Cas9 and Cas12 respectively. For heterozygous SNPs not directly covered, we also provide the nearest allele-specific target within a 500pb window. As additional genome editing systems are adopted for use in *C. albicans*, the coverage of heterozygous sites for LOH engineering will be increased, such that almost all sites could be targeted for CRISPR-induced LOH.

## Discussion

In *C. albicans*, LOH events are commonly observed in evolved strains or clinical isolates, but their potential contribution to resistance is rarely measured because of the complexity of reproducing these structural variants in the lab. Here, we developed an approach that uses the CRISPR-Cas9 system and allele-specific sgRNAs to streamline LOH engineering. Our method is efficient enough so that no LOH-associated phenotype is required to enrich for edited clones. Most recombination tracts are relatively short, similar to naturally occurring events. We show that this approach can be used to reproduce LOH events involved in resistance and that most heterozygous sites in the genome can be targeted directly or indirectly via a neighboring site.

While they generally exhibit lower variant density, many pathogenic species closely related to *C. albicans* are also heterozygous diploids. In *Candida parapsilosis*, CRISPR-Cas9 genome editing has been known to induce off-target LOH events near the target site to a greater extent compared to classical recombination approaches (Lombardi *et al*. 2022). Recombination tracts were much longer than what we observed in our edited strains, but the sgRNAs used were not allele-specific, which might have resulted in more cut and repair cycles before editing stopped. Even if unintentional, this observation suggests that allele-specific sgRNAs might also allow LOH engineering in other fungal pathogens. It is known that CRISPR-Cas9 editing in *C. albicans* is inefficient if a repair template is not provided (Vyas *et al*. 2015). Similarly, CRISPR-Cas9 targeting without a recombination template is also extremely inefficient in *S. cerevisiae* (DiCarlo *et al*. 2013), suggesting similar biases in how DNA repair is handled after the double-stranded break is made. Whether another species’ DNA repair processes after double-stranded breaks match this pattern might be the determining factor that will influence the applicability of this method in other heterozygous diploid fungal species. The code we used to generate the *C. albicans* sgRNA (https://github.com/TheShapiroLab/CRISPR_LOH/) databases has been designed to be modular and easy to convert for use with other nucleases and species, with the caveat that it requires a high-quality phased genome assembly for use.

Although it can efficiently induce LOH events, our method still provides limited control over the exact length of the resulting recombination tracts. In scenarios where the goal would be to generate many different events to map out precisely what parts of a LOH are driving a phenotype of interest, this is an advantage. A single CRISPR transformation can allow for the generation of multiple strains with different independent events, which can then be isolated via re-streaking steps to avoid issues with colony heterogeneity. Conversely, the addition of tailored donor DNA as a recombination template could help increase both editing rates and outcome purity in cases where the goal is to reproduce a specific LOH event. In *C. albicans,* long repeats within the genome have been identified as potential hotspots for recombination in LOH events (Todd *et al*. 2019). In the case of our *HIS4* and *KSR1* LOH events, the end of the recombination tracts did not match an annotated repeat, suggesting they did not influence editing outcomes. This is encouraging, as it means these repeats are not necessarily limiting factors in the range of possible edits.

Examining the genomes of cells that have gained a phenotype of interest, like antifungal resistance, is often the first step to understanding the molecular mechanisms behind it. Once identified, candidate driver mutations should ideally be recreated and validated in the lab. In the case of antifungal drug resistance, these validations are rarely done (Bédard *et al*. 2024a), especially in the case of more complex mutations like LOH. Our method will therefore facilitate the direct characterization of this understudied class of mutational events that can play a key role in the emergence of antifungal resistance.

## Materials and Methods

### Media and growth conditions

5-alpha Competent *Escherichia coli* cells from NEB (cat. C2987H) were grown at 30°C in Lysogeny Broth (LB) and LB plates supplemented with both 100μg/mL ampicillin (AMP) and 250μg/mL nourseothricin (NAT) for plasmid selection. SC5314 *Candida albicans* cells were grown routinely at either 30°C or 37°C in yeast peptone dextrose (YPD) broth and YPD plates supplemented with 250μg/mL NAT for plasmid selection.

### sgRNA cloning

The plasmid backbone used in this study was our *C. albicans* active CRISPR-Cas9 plasmid (pRS118), which we have now deposited to Addgene (plasmid #234878) (Bédard *et al*. 2024b). Each sgRNA was ordered in both a forward and reverse-complement orientation from Integrated DNA Technologies (IDT) and cloned into pRS118 (plasmids: Table S3, oligonucleotides Table S4) as previously described (Gervais *et al*. 2023). Briefly, the two complementary oligos for each sgRNA were heated separately at 94°C for one minute, mixed together in equal concentrations, and heated again at 94°C for two minutes before leaving to cool at room temperature. The duplexed oligo was then cloned into pRS118 via a Golden Gate strategy with ∼1000ng of plasmid along with the following: 1µL of duplexed oligo, 2µL of 10X rCutSmart buffer, 2µL of ATP, 1µL of SapI, 1µL of T4 DNA ligase, and raised to a total volume of 20µL with nuclease-free water. Each mixture was incubated in a thermocycler using the following cycling conditions: (37°C for 2 min, and 16°C for 5 min) for 99 cycles; 65°C for 15 min; 80°C for 15 min. Afterwards, an additional 1µL of SapI was added to each reaction and incubated at 37°C for 1h to remove any plasmids not containing the properly integrated duplexed oligo fragment. Then, 3µL of each mixture was transformed into 5-alpha Competent *Escherichia coli* cells from NEB (cat. C2987H), plated onto LB media containing AMP and NAT, and left to grow at 30°C static for 1-2 days. Transformants were PCR tested for the presence of the corresponding sgRNA as previously described before proceeding (Wensing and Shapiro 2022). Plasmids were miniprepped with a GeneJET Plasmid Miniprep Kit from ThermoFisher (cat. K0503) as per the manufacturer’s instructions.

### *Candida albicans* transformations

*C. albicans* transformations were performed as previously described with some minor modifications (Wensing and Shapiro 2022). *C. albicans* cells were grown up overnight in YPD at 25°C (250 RPM) to help ensure the cultures were still in growth phase the following morning. Plasmids were linearized using the PacI enzyme from NEB (cat .R0547L), and left at 37°C for 12h followed by a 20 min heat-inactivation step at 65°C. A transformation master mix was prepared with 800µL of 50% polyethylene glycol (PEG), 100µL of 10X Tris-EDTA buffer solution, 100µL of 1M lithium acetate, 40µL of UltraPure™ Salmon Sperm DNA Solution from ThermoFisher (cat. 15632-011), and 20µL of 1M dithiothreitol (DTT). The transformation mix was added to pelleted *C. albicans* cells, followed by the addition of the linearized CRISPR plasmid, and left to incubate at 30°C for 1h. The solutions were then heat shocked at 42°C for 50 min. Pelleted heat-shocked cells were washed with fresh YPD three times, then grown in YPD for 4h to allow for expression of the NATr construct. Transformed cells were then plated on YPD containing NAT, grown at 30°C for 2 days, and transformants were picked haphazardly for phenotypic assays. For the experiment using donor DNA, oligonucleotides oRS1632 and oRS1633 were used to amplify a ∼1350 bp region centered on the target of pRS893 from genomic DNA of strain AA_4, and ∼200 ng of this product was added to the transformation mix.

### Spot Plating

Transformants were individually picked and diluted into 80uL of YPD on a 96-well plate. Then, 5µL of each well was spotted onto both solid synthetic complete (SC) and SC-His dropout media (CSM-His Powder, 10 grams from Sunrise Science Products, cat. 1006-010) and briefly left to dry. The plates were left to grow at 30°C overnight, and *HIS4* LOH was determined the next day by the presence of growth on SC and the absence of growth on SC-His.

### Minimum Inhibitory Concentration (MIC) Assays

Fluconazole MIC assays were performed in 96-well flat-bottomed plates. A suspension of 128μg/mL fluconazole was prepared in YPD, and 200μL was added to each well in the first column of the plate. The drug was then serially diluted ½ across each subsequent column of the plate in YPD, until the second-to-last column. Overnight cultures of *C. albicans* grown in YPD at 30°C (250 RPM) were then diluted to an OD600 of 0.1 in 1mL of YPD, and 100μL of this suspension was then further diluted into 6mL of YPD. 100μL of each diluted culture was then added to one row each of the 96-well plate, for a final OD600 of ∼0.000833. The gradient of fluconazole, ranged from 64μg/mL to 0μg/mL. Each plate contained one row of WT SC5314 *C. albicans* as a control, and a row of blank media as a contamination control. All isolates were tested in duplicate to determine the presence of the drug-resistant phenotype, and plates were incubated at 37°C (static) for 24h. OD600 values were read using an Infinite 200 PRO microplate reader (Tecan).

### gDNA Extraction

Genomic DNA was prepared using a PureLink™ Genomic DNA Mini Kit from Thermo Fisher Scientific (cat. K182001) broadly as per the manufacturer’s instructions. Briefly, strains were grown up overnight in 5mL of YPD (250 RPM) at 30°C. The next morning, 1mL aliquots were frozen at −80°C, thawed, and pelleted at 13000 x G for 1 min. Pellets were reuspended in 600μL of homemade sorbitol buffer, which was prepared in nuclease-free water with 1M sorbitol, 0.1% 2-mercaptoethanol, 0.01M EDTA, and ∼200 units of zymolase from BioShop (cat. ZYM001.1) per reaction. Then, the resuspended pellets were left at 35°C (static) for 2h to generate spheroplasts. Spheroplasts were then pelleted at 4000 x G for 10 minutes, resuspended in 180μL of PureLink™ Genomic Digestion buffer and 20μL of Proteinase K, and incubated at 55°C (static) for 45 minutes. 20μL of RNase A (20 mg/mL) was then added to the lysates, samples were incubated at room temperature for a few minutes, and then 200μL of PureLink™ Genomic Lysis/Binding Buffer and 200μL of 100% EtOH was added. Samples were then transferred to a PureLink™ Spin Column and washed/isolated as per the manufacturer’s instructions.

### Nanopore Sequencing and Genome Analyses

All LOH colonies sequenced were from independent transformations to ensure each population originated from a different editing event. For the histidine auxotrophs, the target gene in selected colonies was first sequenced via Sanger to confirm the presence of the LOH, using primers available in Table S4. Yeast Genome Sequencing was performed by Plasmidsaurus using Oxford Nanopore Technology with custom analysis and annotation. Nanopore reads were aligned to the ASM18296v3 SC5314 haploid reference genome (Skrzypek *et al*. 2017) using minimap2 version 2.28 (Li 2018), and the alignments were indexed using samtools v1.20 (Li *et al*. 2009). The average coverage across the genome for each strain was calculated using the `genomecov` function of Bedtools v2.31 (Quinlan and Hall 2010). The phased diploid reference genome, genomic feature annotations, and the list of heterozygous sites were downloaded from the Candida Genome Database (CGD, (Skrzypek *et al*. 2017)). Heterozygous sites around the targeted sites were examined using pileups generated by `samtools pileup` and manually inspected using IGV (Robinson *et al*. 2011). LOH tract length was calculated as the length of DNA from the first to the last heterozygous site detected on each side of the sgRNA target.

### Allele-specific sgRNA database design

We used a custom Python script to scan the reference genome for Cas9 or Cas12a target sites using regular expressions. Potential sites were then filtered to 1) remove sgRNAs with homopolymer repeats (n>4) and 2) filter for sequences only present in one of the two chromosomes. These sites were then exported to a FASTA file and aligned on the diploid reference genome using Bowtie 2 (Langmead and Salzberg 2012). The proportions of sgRNAs targeting coding or non-coding genomic features were computed using featureCounts (Liao *et al*. 2014). We cross-referenced sgRNA read density generated with samtools mpileup with the set of known heterozygous sites to compute coverage metrics for SNPs.

## Data Availability Statement

Strains and plasmids are available upon request. Raw sequencing data files have been deposited on the NCBI SRA under PRJNA1260133 (release pending). The code used for data analysis is available at (https://github.com/TheShapiroLab/CRISPR_LOH/). The database of allele-specific sgRNAs is provided as Tables S5 and S6.

## Acknowledgements

The authors thank M. Hénault for helpful comments on the manuscript. This work was supported by the FRQS through a postdoctoral fellowship to PCD, by NSERC through a CGS D. award to NCG, CIHR through grant PJT 162195 to RSS, CIFAR through a grant from the Fungal Kingdom: Threats & Opportunities program to both CAC and RSS, and NIAID through grant U19 AI110818. RSS holds the Canada Research Chair in Microbial Functional Genomics and Synthetic Biology.

## Supplementary Figures 1-3

**Figure S1:**
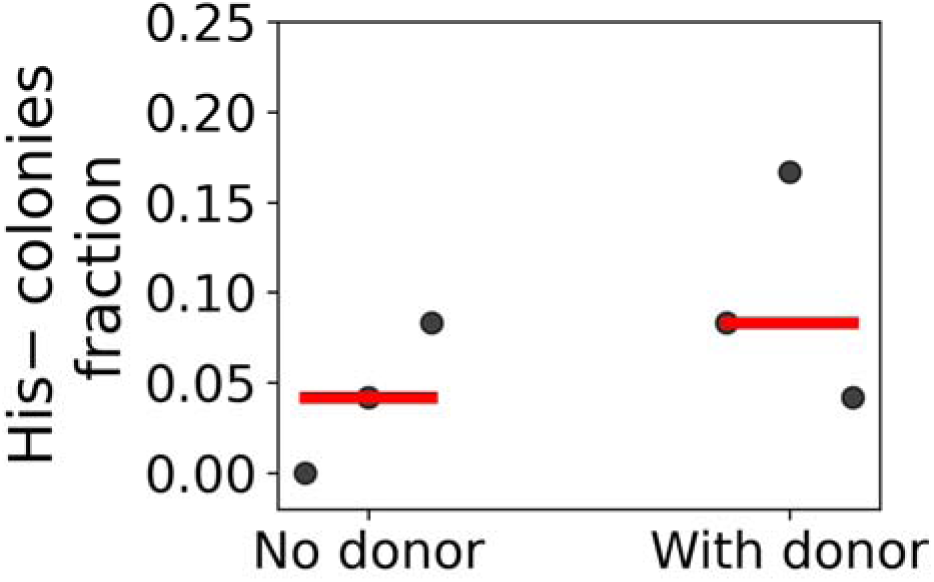
Adding donor DNA to the transformation mix improves LOH editing efficiency. Donor DNA generated from a strain with the LOH of interest (fRS1224) was added to the transformation mix.

**Figure S2:**
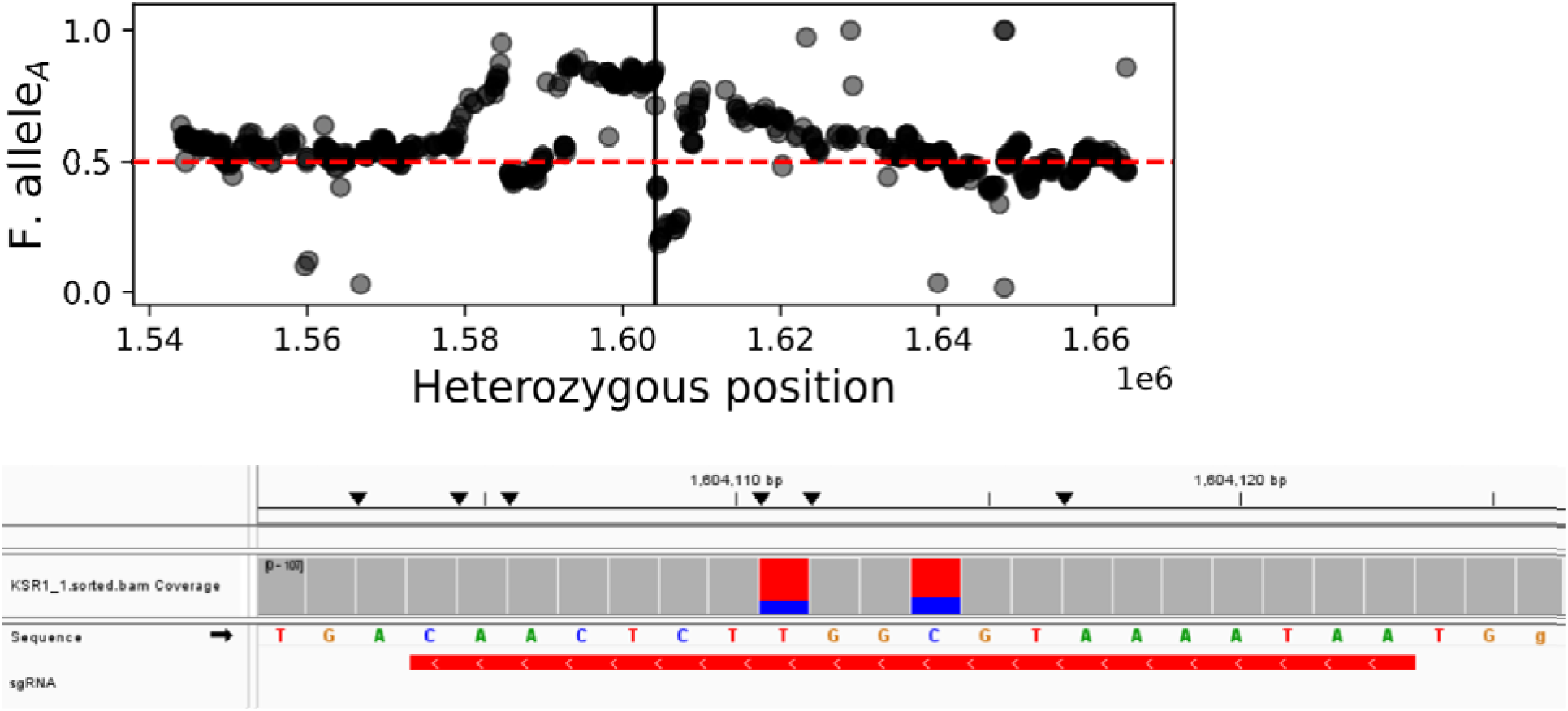
KSR1_1 does not have the desired LOH edit. Chromosome A allele frequency shows complex patterns indicative of multiple LOH events from A to B as well as from B to A. A de novo mutation (C>T) is also present at the sgRNA target site.

**Figure S3:**
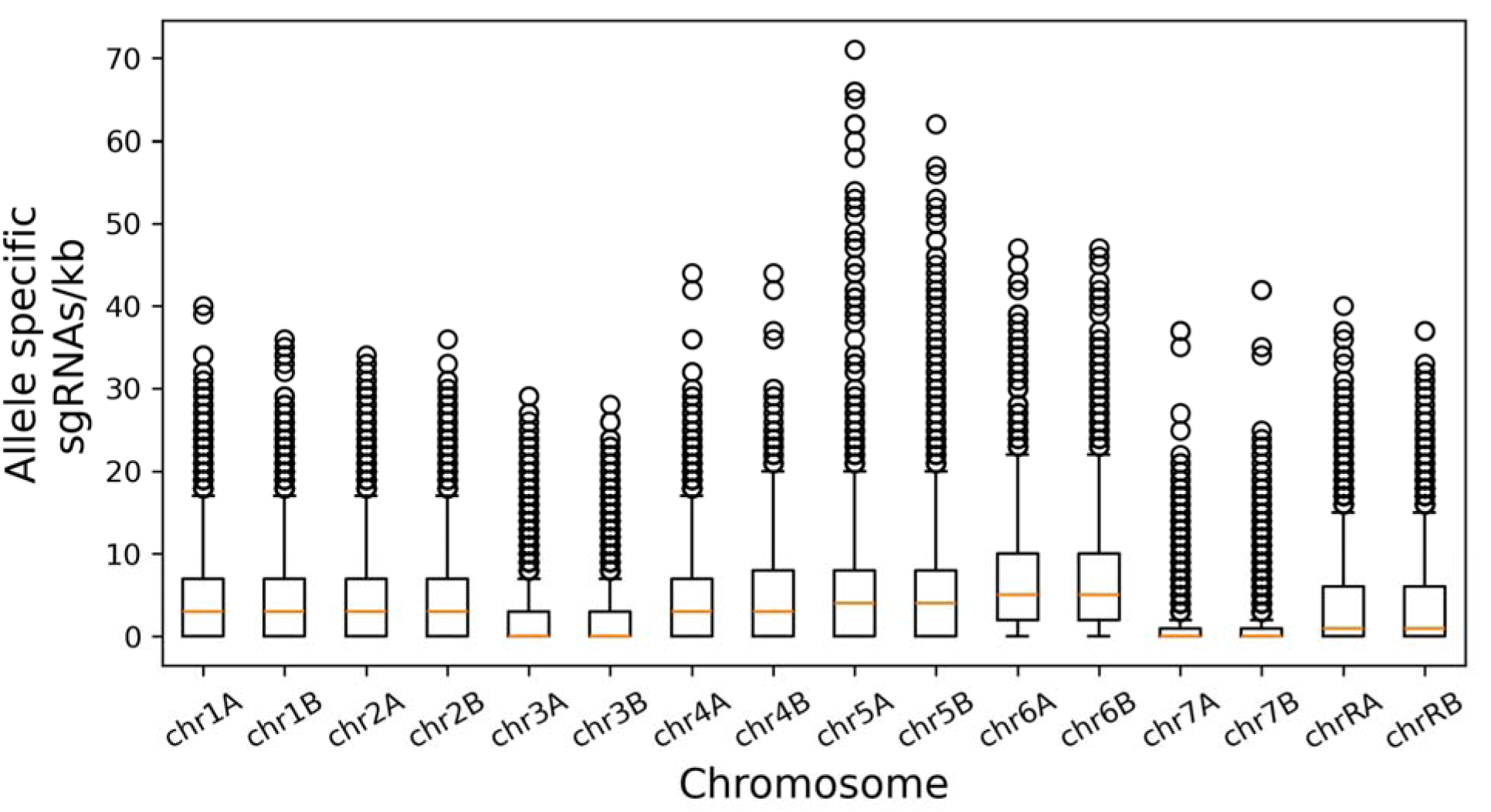
Cas9 allele target density is high across large parts of the *C. albicans* genome. The number of sgRNAs specifically targeting either the A or the B chromosome was calculated using rolling 1kb windows for the entire genome. Chromosomes 3, 7 and R are known to harbor large LOH tracts in SC5314, resulting in fewer heterozygous sites to target.

